# A more physiological approach to lipid metabolism alterations in cancer: CRC-like organoids assessment

**DOI:** 10.1101/462929

**Authors:** Silvia Cruz-Gil, Ruth Sanchez-Martinez, Sonia Wagner-Reguero, Daniel Stange, Sebastian Schölch, Kristin Werner, Ana Ramirez de Molina

## Abstract

Precision medicine might be the response to the recent questioning of the use of metformin as an anticancer drug in colorectal cancer (CRC). Thus, in order to establish properly its benefits, its application need to be assayed on the different progression stages of CRC. In this way, organoids imply a more physiological tool, representing a new therapeutic opportunity for CRC personalized treatment to assay tumor stage-dependent drugs effects. Since the lipid metabolism-related axis, ACSL/SCD, stimulates colon cancer progression and Metformin is able to rescuing the invasive and migratory phenotype conferred to cancer cells upon this axis overexpression; we checked ACSL/SCD status, its regulatory miRNAs and the effect of Metformin treatment in organoids as a model for specific and personalized treatment. Despite ACSL4 expression is upregulated in CRC-like organoids, Metformin is able to downregulate it, especially in the first stages. Besides, organoids are clearly more sensitive in this first stage (Apc mutated) to Metformin than current chemotherapeutic drugs such as fluorouracil (5-FU). Metformin performs an independent “Warburg effect” blockade to cancer progression and is able to reduce crypt stem cell markers expression such as Lgr5+. These results suggest a putative increased efficiency of the use of Metformin in the first stages of CRC than in advanced disease.

## Introduction

Colorectal cancer (CRC) is the third most common cancer in men (10% of the total), after lung and prostate cancer, and the second in women (9.2% of the total), after breast cancer [1]. Most of the CRC cases are sporadic (70-80%), which consists of the acquisition of somatic mutations and in which there is no family history or genetic predisposition. The remaining cases (20-30%) are those among close relatives, which are divided into inherited or familial CRC, [2]. Genetically, sporadic CRC development is due to the abnormalities accumulation in tumor suppressor genes and oncogenes [3,3]. Previous research postulated the adenoma-carcinoma transition theory, in which specific somatic mutations promoting tumorigenesis are acquired; proposed by Fearon and Vogelstein (Vogelgram). The Vogelgram proposes that the adenoma-carcinoma sequence model would start with loss of the *APC* gene, followed by mutations in *KRAS* or *BRAF* genes, mutations or loss of *TP53* gene and of SMAD family member 4 (*SMAD4*) [4].

Over the last decade, the interest in metabolic research with respect to cancer has been expansively increased. The first and most characterized tumor metabolism event to be described is the exacerbated glucose uptake and glycolysis utilization; which even in normoxic condition, are not used for maximal ATP generation via mitochondrial respiration. This phenomenon is denoted as the “Warburg effect”.

Even though lipid-associated pathways are functionally dependent on glucose and glutamine catabolic pathways, are now a well-recognized and frequently described cancer metabolic feature with a key role in their tumorigenesis. This is the case for the ACSL/SCD axis [5], a lipid metabolism-related network described to promote tumorigenesis through an epithelial-mesenchymal transition (EMT) program that promotes migration and invasion of colon cancer cells. The mesenchymal phenotype produced upon overexpression of these enzymes is reverted through reactivation of AMPK signaling performed by the well-known anti-diabetic drug, Metformin. Though its mechanism of action is not fully understood, Metformin has shown a robust anti-proliferative effect on several types of cancer such as colon, pancreatic, breast, ovarian, prostate and lung cancer cells [6]. Furthermore, Metformin has been recently associated with improved survival of cancer patients, including CRC, though its use as an antitumoral agent has not been established yet [7]

The ACSL/SCD axis pro-tumorigenic activity has been also described to be post-transcriptionally regulated by miRNAs. miR-544a, miR-142, and miR-19b-1 has been proposed as major regulators of the ACSL/SCD network and the miR-19b-1-3p isoform decreased expression associated with a poorer survival rate in CRC patients, consistently with ACSL/SCD involvement in patients relapse [8].

To get insight into the metabolic implication on CRC progression with a special focus on the ACSL/SCD axis and the effect of metformin in each case, more personalized and physiological tools are needed since most of the available data rely on traditional studies using cancer cell lines cultures. In this way, the organoid culture system opens a new methodological door for *ex vivo* studies.

Adult tissue-derived epithelial organoids, also called “mini guts” [9] are stereotypic tissue-like structures derived from digestive healthy tissues or tumors which mimics *in vitro* the tissue composition and morphology of their *in vivo* counterparts [10]. This methodology was first established in long-term primary culture from mouse small intestinal crypts to generate epithelial organoids with crypt- and villus-like epithelial domains representing both progenitor and differentiated cells [11].

The organoids technology takes advantage of the intestinal epithelium self-renewing capacity. Organoids starts from LGR5+ gut epithelial stem cells forming symmetric cyst structures, which finally will form budding structures resembling intestinal crypts. These budding structures are formed by these LGR5+ stem cells flanked by differentiated daughter cells [9].

Organoids are currently employed in colorectal cancer studies and chemotherapy assessment [12,13]. Along with intestinal organoids, similar epithelial organoids culture conditions for other mouse and human digestive epithelial tissues have been also adapted [14–17] including tumor-derived organoids from cancer patients. Importantly, organoids grow as pure epithelial cultures without any contamination of vessels, immune cells or non-transformed mesenchymal which leads to an accurate sequencing or expression profiling [10].

## Materials and Methods

### CRC-like organoids: culture and maintenance

#### Mice

Mutant intestinal murine organoids were obtained from the Universitätsklinikum Carl Gustav Carus, Dresden. All procedures involving animals were conducted strictly in accordance with FELASA regulations and approved by the animal welfare committees of the Technische Universität Dresden and the Landesdirektion Sachsen prior to initiation of the experiments.

Mice with conditional mutations in Apc, Kras, Tp53 and Smad4 were obtained from the NCI Mouse Repository (Apc, Kras Tp53) or the Jackson Laboratory (Smad4) and interbred to obtain compound mutant mice (Table 1). The CRC-like organoid model represents the adenoma-carcinoma sequence with the most common acquired mutations in a sporadic CRC: *APC*^fl/fl^*, KRAS*^G12D/WT^*, P53*^R172H/WT^ and Smad4^fl/fl^ (corresponding to stages I to IV) (Table 2). The parental mouse lines were described in Table 2.

**Table 1:** Parental mouse lines and publication’s PMID of the mutations in the organoids

| Mutation | Designation | PMID |
| --- | --- | --- |
| *APC* | NCI: 01XAA | 17002498 |
| *KRAS*<sup>G12D</sup> | JAX: 008179 | 11323676 |
| *P53*<sup>R172H</sup> | JAX: 008652 | 15607980 |
| *P53 floxed* | NCI: 01XC2 | 11694875 |
| *Smad4 floxed* | JAX: 017462 | 11857783 |

**Table 2:**
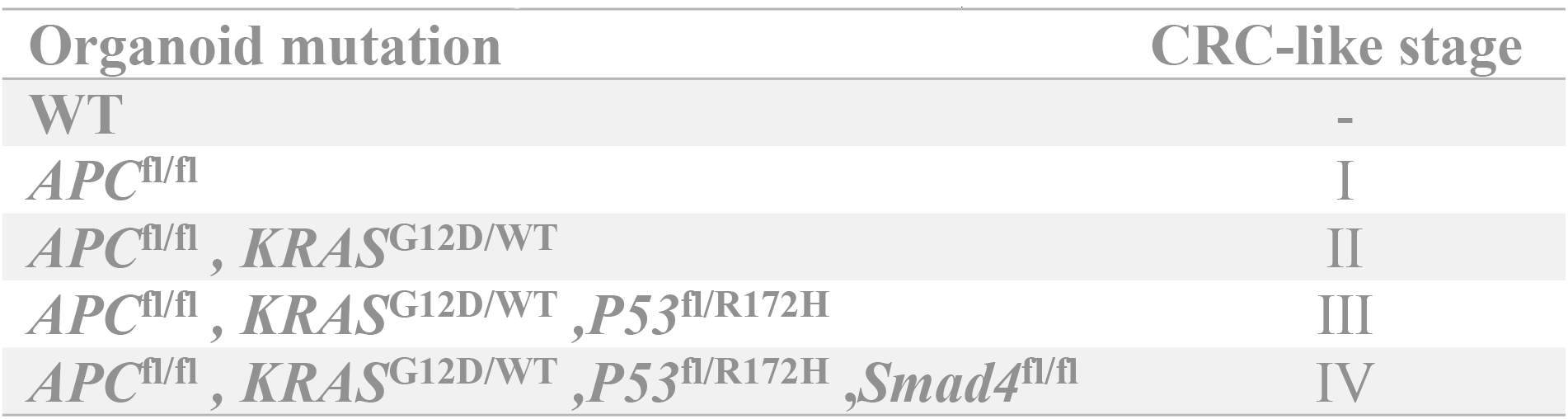
CRC-like organoids with the acquired mutations related to the stage.

| Organoid mutation | CRC-like stage |
| --- | --- |
| WT | - |
| *APC*<sup>fl/fl</sup> | I |
| *APC*<sup>fl/fl</sup>, *KRAS*<sup>G12D/WT</sup> | II |
| *APC*<sup>fl/fl</sup>, *KRAS*<sup>G12D/WT</sup>, *P53*<sup>fl/R172H</sup> | III |
| *APC*<sup>fl/fl</sup>, *KRAS*<sup>G12D/WT</sup>, *P53*<sup>fl/R172H</sup>, *Smad4*<sup>fl/fl</sup> | IV |

Murine organoids mutagenesis is conditioned by the Cre/loxP system. Adenoviral infections were performed as explained in [18] to provide active mutations.

#### Crypt isolation and organoid culture

Crypts were isolated from the murine small intestine by incubation for 30 min at 4°C in PBS containing 2 mM EDTA as previously reported [11,19]. Isolated crypts were seeded in Matrigel (Corning^®^ Matrigel^®^ Matrix). The basic culture medium (Advanced Dulbecco’s modified Eagle Medium DMEM/F12 complemented with penicillin/streptomycin, 10 mmol/L HEPES, 1x Glutamax [Gibco], named ADF +++) was supplemented with: 100 ng/ml Noggin (Peprotech), R-spondin (conditioned medium, 10% final volume), 1x B27 (Invitrogen), 1x N2 (Invitrogen), 1,25 mM N-acetylcysteine (Sigma-Aldrich), 100 μg/mL Primocin TM (InvivoGen) and 50 ng/mL mEGF (Thermofisher). The complete media is named supplemented ADF +++ media. For passaging, organoids were removed from Matrigel and mechanically dissociated with a glass pipette, pelleted and then transferred to fresh Matrigel [11,14,20]. Splitting was performed twice a week in a 1:3 split ratio. Cultures were kept at 37 °C, 5% CO2 in humidity.

### Drugs treatment - viability assays

Cell viability was determined by counting and seeding 1000 crypts in 60% of Matrigel in 48-well plates. After 2 days of culture, organoids were exposed 48 hours to 10 μM Metformin (Sigma) or 10, 100 or 150 μM 5-FU (Sigma) in supplemented ADF +++ media, as indicated in the figures. At this point, organoids were collected, split and reseeded for recovery experiments over 72 hours in supplemented ADF +++ media.

Upon treatments (48h) or recovery assays (post-72h), organoids were incubated 3 hours with 3- (4,5-dimethyl-thyazol-2-yl)-2,5-diphenyl-tetrazolium (MTT, Sigma). After discarding the media, 20 μl of 2% SDS (Sigma) solution in H2O was added to solubilize Matrigel (2 h, 37 °C). The resultant formazan was dissolved in 100 μl of DMSO for 1 h (37 °C). The absorbance was measured on the microplate reader (Asys UVM 340, Isogen life science) at 562 nm.

Untreated organoids were defined as 100% viable. Data were expressed as the fold change of viable cells from treated organoids compared to the non-treated organoids.

### RNA isolation and RT-QPCR

For RNA isolation, organoids were released from Matrigel (Corning) with cold Dispase (Corning) and pelleted by centrifugation. The supernatant was removed and pelleted organoids were carefully resuspended in Trizol (Qiagen), and storage at −80°C. RNA was isolated according to the supplier’s protocol (Invitrogen) and the concentration and purity (A260/A280 ratio) were determined by spectrophotometric analysis (NanoDrop 2000 Spectrophotometer ThermoScientific). 20 ng/μl RNA was reverse-transcribed using the High Capacity RNA-to-cDNA kit (ThermoFisher), according to manufacturer’s instructions. Relative gene expression was measured using VeriQuest Fast SYBR Green qPCR Master Mix (2X) (Isogen). Primers used are listed in S1 Table. Regarding miRNAs, their expression was monitored using TaqMan^®^ MicroRNA Reverse Transcription Kit (ThermoFisher Scientific) and Taq-man miRNA probes for RT-qPCR (S2 Table). RT-QPCRs were performed on the QuantStudio 12K Flex (Applied Biosystems) and the 2^-ΔΔCt^ method was applied to calculate the relative gene or miRNA expression.

### L-Lactate quantification

Organoids were seeded at a density of 1000 crypts per well in a 48-well plate. After 48 hours, the medium was changed to PBS, 10 mM of Metformin or 10 μM 5-FU in supplemented ADF +++ media overnight at 37°C before quantification. Using Cayman’s Glycolysis cell-based assay (Cayman, Ann Arbor, MI, USA, 600450) extracellular L-Lactate was measured by determining absorbance at 490 nm. L-Lactate measurements (mM) were normalized to total protein concentration (mg) x100.

### Statistical analysis

All statistical analyses were performed using the Graph Pad Prism software (Ver. 7.03) (GraphPad Software, San Diego, CA, USA). Significance between groups was determined by *t*-test analyses (unpaired Student’s t-tests). Data with *P* < 0.05 were considered statistically significant (ns, *P* > 0.05; *, *P* ≤ 0.05; **, *P* ≤ 0.01; ***, *P* ≤ 0.001; ****, *P* ≤ 0.0001). All reported *p* values were two-sided. All values are reported as mean ± S.D.

## Results

### ACSL4 is overexpressed throughout CRC-like organoids stages

ACSL4 has been previously reported to be overexpressed in malignant tumors, and together with ACSL1 and SCD form an axis involved in CRC progression. ACSL1, ACSL4, and SCD mRNA expression was measured in CRC-like organoids. ACSL4 mRNA was very significantly augmented in more aggressive stages compared to WT (Fig 1A). It is shown an intermediate expression pattern in Apc-mutated organoids, with a significantly differential expression (*p-value:* **) compared to the following second stage (Apc, Kras mutated stage) henceforth. Conversely, ACSL1 and SCD levels were maintained or increased from the third stage henceforth, respectively (S1 Fig).

**Figure 1:**
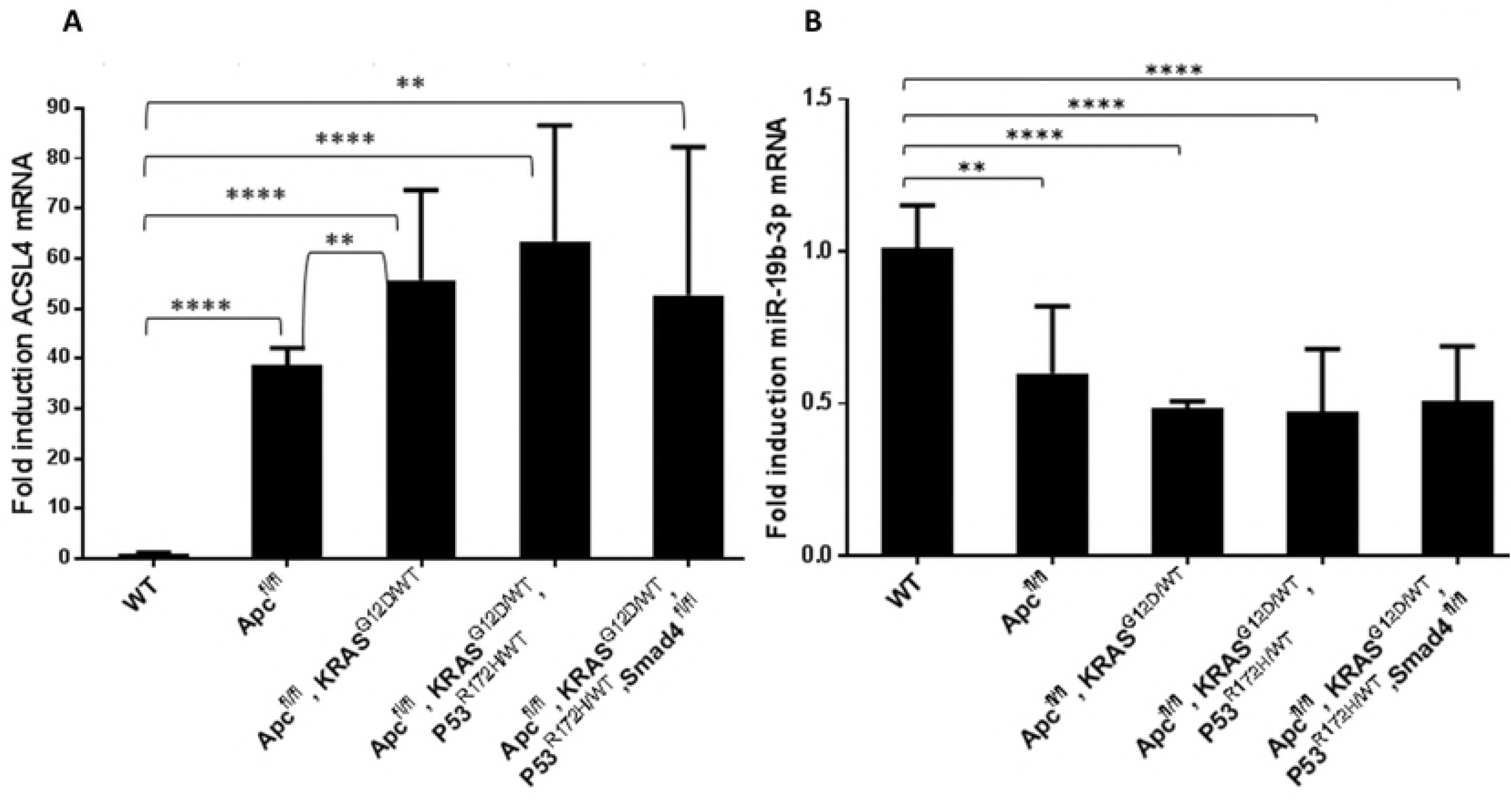
ACSL4 is overexpressed throughout CRC-like organoids stages while miR-19b-1- 3p preserves its protective role. A) RT-QPCR analysis showing ACSL4 mRNA expression levels throughout CRC-like organoids stages. B) RT-QPCR analysis showing miR-19b-1-3p mRNA expression levels throughout CRC-like organoids stages. Results represent the fold-change mean ±SD (*n* = 4) in plots A, (*n* = 3) in plots B. (ns, *P* > 0.05; *, *P* ≤ 0.05; **, *P* ≤ 0.01; ***, *P* ≤ 0.001; ****, *P* ≤ 0.0001)

Interestingly, organoids in more advanced stages (III and IV) presented a genetic misbalance in ACSL4 expression (Fig 1A) with huge differences in their fold inductions ranges in the same stage, though with a similar tendency.

### MiR-19b-1-3p keeps its protective role in CRC-like organoids

MiRNAs expression was assayed in 3 different RNA extractions over time. Previous results from our group pointed toward a correlation between miR-19b-1-3p lower expression and a poorer prognosis in CRC patients (which might have a putative high clinical interest due to its potential to be assed in plasma as a non-invasive biomarker); very likely through its involvement in cell invasion and lipid metabolism regulation [8]. In the case of CRC-like organoids, this tendency was maintained and miR-19b-1-3p expression was decreased in a stage-dependent manner (Fig 1B).

Together with miR-19b-1-3p, miR-142 (3p and 5p isoforms) and miR-544a (without murine isoform) were also involved on targeting ACSL/SCD axis [8]. Hence, the previous mentioned miRs plus miR-19b-1-5p isoform was measured though no statistically significant differences were found in its expression (S2 Fig)

### Metformin decreases CRC-like organoids viability to the same extent as current chemotherapy without significant effects on WT organoids

Since Metformin treatment, an AMPK activator used as antidiabetic treatment that has been recently associated to increased survival of cancer patients, was able to rescue the epithelial phenotype from the EMT process caused by the overexpression of ACSL/SCD in CRC cells [5]; we wondered what this drug effect would be through the different stages in tumor progression. CRC-like organoids were treated with PBS, 10 μM of Metformin or with the commonly used chemotherapeutic agent 5-FU; and the organoids viability was examined by MTT assays 48 hours upon treatment. None of the drugs affected significantly the viability of WT organoids (Fig 2A), while they were able to cause a decrease of about 50% in the viability of mutated organoids corresponding to the most aggressive phenotypes (Fig 2B-E). 5-FU higher concentrations (100μM and 150 μM) showed the same effects than the lower concentration (10 μM) in mutated organoids, while they had stronger effects on WT ones (S3A-E Figs).

**Figure 2:**
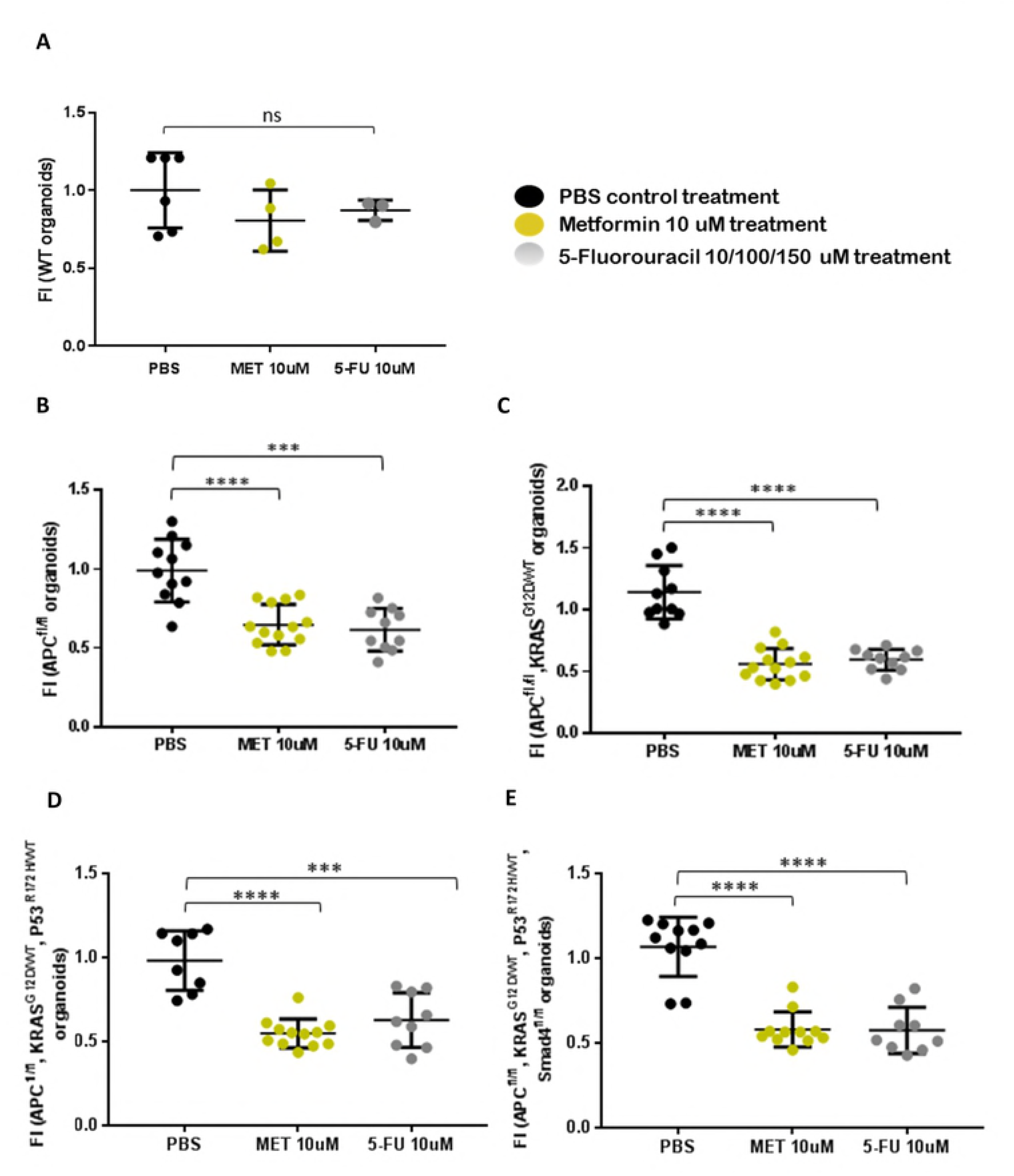
Metformin decreases CRC-like organoids viability to the same extent as current chemotherapy without significant effects on WT organoids. MTT cell viability assays upon 48 hours treatments with Metformin or 5-FU in the different CRC-like organoids representative stages (A) WT organoids; (B) *APC*^fl/fl^ organoids resembling stage I; (C) *APC*^fl/fl^ *, KRAS*^G12D/WT^ organoids resembling stage II; (D) *APC*^fl/fl^ *, KRAS*^G12D/WT^ *,P53*^R172H/WT^ organoids resembling stage III; (E) *APC*^fl/fl^ *, KRAS*^G12D/WT^ *,P53*^R172H/WT^, Smad4^fl/fl^ organoids resembling stage IV. Data are represented by the fold-change mean ±SD (*n* = 3) in all the plots except A: (n=2). (ns, *P* > 0.05; *, *P* ≤ 0.05; **, *P* ≤ 0.01; ***, *P* ≤ 0.001; ****, *P* ≤ 0.0001).

### Metformin treatment recovery is significantly lower compared to 5FU in first stages organoids while WT organoids present an opposite behavior

To further check the treatments scope, and analyzing not only the effect but also the potential reversibility of the treatment in normal and tumoral cells in different stages, organoids viability was assayed upon 48 hours treatment (PBS, Metformin or 5-FU) plus the subsequent recovery of 72 additional hours in their growing media. In this case, WT organoids showed differential recovery sensitivity to the treatment. Metformin treated and recovered WT organoids presented almost similar measurements than only treated organoids. Nonetheless, 5FU treated WT organoids recoveries are noteworthy more sensitive and upon 72h recovery time their viability was quite significant reduced (*p-value:* ***) (Fig 3A). In Apc mutated organoids the recovery is very significantly lower upon Metformin treatment (*p-value:* ****) than 5-FU (*p-value:* *), compared to PBS recovery control; making these Apc mutated organoids the most responsive to the Metformin treatment compared to 5-FU (Fig 3B). Regarding organoids corresponding to stages II-III (Figs 3C and D), both treatments presented almost similar recovery effects, while in stage IV organoids, 5-FU presented a stronger effect shown by the lower recovery of these 5-FU treated organoids (Fig 3E). Again 5-FU higher concentrations (100 μM and 150 μM) had nearly the same recovery effects than the lower concentration (10 μM) (S4A-E Figs).

**Figure 3:**
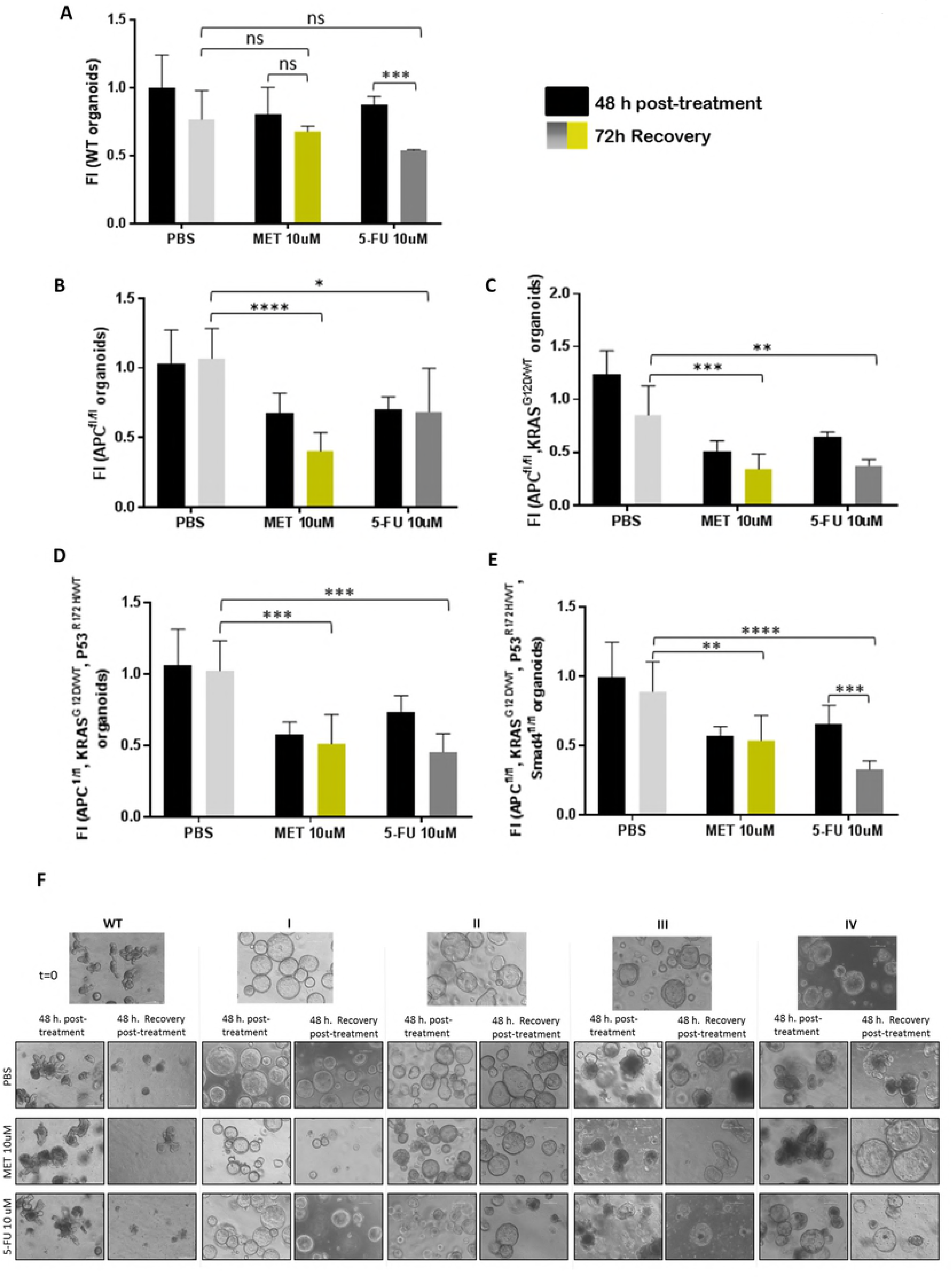
Metformin treatment recovery is significantly lower compared to 5FU in first stages organoids while WT organoids present an opposite behavior. MTT cell viability assays upon 48 hours treatments (black bars) and upon extra 72h post-treatment recovery with PBS (light grey bars) Metformin (yellow bars) or 5-FU (dark grey bars) in the different CRC-like organoids representative stages (A) WT organoids; (B) *APC*^fl/fl^ organoids resembling stage I; (C) *APC*^fl/fl^ *, KRAS*^G12D/WT^ organoids resembling stage II; (D) *APC*^fl/fl^ *, KRAS*^G12D/WT^ *,P53*^R172H/WT^ organoids resembling stage III; (E) *APC*^fl/fl^ *, KRAS*^G12D/WT^ *,P53*^R172H/WT^, Smad4^fl/fl^ organoids resembling stage IV. Data are represented by the fold-change mean ±SD (*n* = 3) in all the plots. (ns, *P*> 0.05; *, *P* ≤ 0.05; **, *P* ≤ 0.01; ***, *P* ≤ 0.001; ****, *P* ≤ 0.0001). (F) Organoids pictures with PBS, Metformin or 5-FU, upon 48 hours treatments and upon extra 72h post-treatment recovery as indicated. Pictures were captured using the × 10 objective, in bright field. Leica microscope (Leica microsystems).

Since WT organoids require more time to achieve their size and their crypt-like phenotype (Fig 3F), the recovery measures are lesser than the mutated organoids. On the other hand, WT organoids recovered upon Metformin treatment presented a higher size than the ones treated with 5-FU (Fig 3F). On the contrary, stage I (Apc mutated) organoids recovered upon Met treatment showed an evident reduced size compared with the control and the 5-FU treated ones. In accordance with viability assays results, this effect is lost in further stages; where Metformin is less effective and Metformin treated recovered organoids presented a bigger size than the ones treated with 5-FU.

By way of clarification, all mutated organoids presented Apc mutated since is the first gene in the adenocarcinoma sequence. Apc completed deletion provokes a hyperactive Wnt signaling. This aberration makes an organoids phenotype switch, losing their crypt-like structure and adopting a cystic morphology [17,21]

### Metformin action is stronger on ACSL4 and SCD overexpressing first stages organoids

Since stage I organoids seemed to present a differential sensitivity to metformin compared to other stages, together with a differential expression of ACSL4, we aimed to analyze the possible link between Metformin and ACSL/SCD axis in intestinal organoids.

To this aim, ACSL4 expression was measured, as well as the other enzymes of the ACSL/SCD metabolic network (ACSL1 and SCD) upon 10 μM Metformin treatment. ACSL4 mRNA expression was strongly reduced by this drug compared to their non-treated controls in stage I and II organoids. By contrast, stage III and IV presented no significance in their reduction or a slight significance, respectively. WT organoids also presented a slight reduction of ACSL4 mRNA upon Metformin treatment (Fig 4B). In addition, SCD expression levels were clearly decreased by Metformin in WT and stage I organoids, while a less marked tendency was found for stage III and IV organoids (Fig 4C). ACSL1 mRNA analysis showed less significant results (Fig 4A) upon Metformin treatment.

**Figure 4:**
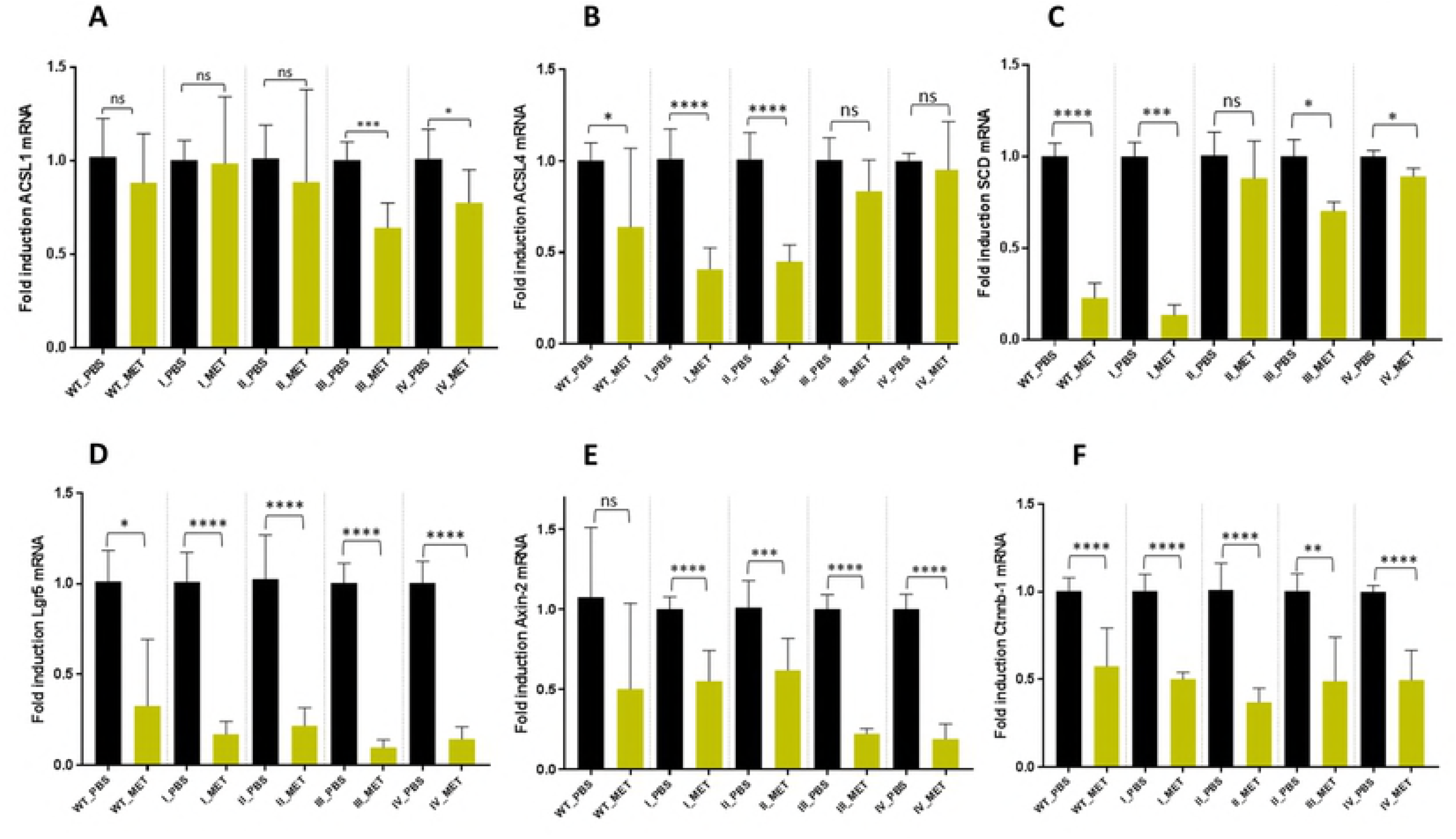
Metformin action is stronger on ACSL4 and SCD overexpressing first stages organoids and downregulates stem cell biomarker LGR5 and Wnt target genes expression in all organoid stages. mRNA expression levels of enzymes related to the ACSL/SCD axis, ACSL4 (A), SCD (B), by RT-QPCR; and expression levels of different stem cell markers, Lgr5 (C), Axin-2 (D) and Ctnnb-1 (E) by RT-QPCR upon PBS (black bars) and 10 uM Metformin (yellow bars). Data are represented by the fold-change mean to each PBS control ±SD (*n* = 3). (ns, *P*> 0.05; *, *P* ≤ 0.05; **, *P* ≤ 0.01; ***, *P* ≤ 0.001; ****, *P* ≤ 0.0001).

The expression of these enzymes was also measured upon 10 μM 5-FU treatment. This drug was able to significantly downregulate ACSL4 and SCD mRNA in most of the stages, though no differences were showed between the effects in initial and later stages such as the case for Metformin treatment (S5A-C Figs).

### Metformin, but not 5-FU, downregulates stem cell biomarker LGR5 and Wnt target genes expression in all organoid stages

To further assay whether Metformin treatment was targeting the organoids crypts stem cell marker, LGR5; we analyzed its expression together with two other Wnt target genes, Axin2 and Ctnnb-1. Importantly, LGR5 expression was significantly diminished in the whole CRC-like organoids series upon Metformin treatment (Fig 4C) as well as Axin-2 (Fig 4D) and Ctnnb-1 (Fig 4E) mRNAs. Surprisingly, this pattern was not maintained when organoids were treated with 10 μM 5-FU (S5D-F Figs).

### Metformin action in CRC-like organoids is not related to a Warburg-effect impairement

The avidity to perform glycolysis even in the presence of oxygen, known as the Warburg effect, is one of the hallmarks of tumors. For this reason, we measured the levels of L-lactate, the end product of glycolysis. CRC-like organoids presented increased glycolysis compared to WT organoids, reflecting an increasing Warburg effect throughout the stages, as expected. Even though 5-FU treatment caused a slight decrease in the glycolytic performance of the mutant organoids (Fig 5), Metformin treatment caused an opposite effect, increasing the glycolytic capacity in all stages, especially in stage I, the most sensitive to the drug. Thus, it seems that Metformin effect on CRC-like organoids viability relies in mechanisms other than preventing pro-tumorigenic Warburg effect, likely through the regulation of lipid metabolism.

**Figure 5:**
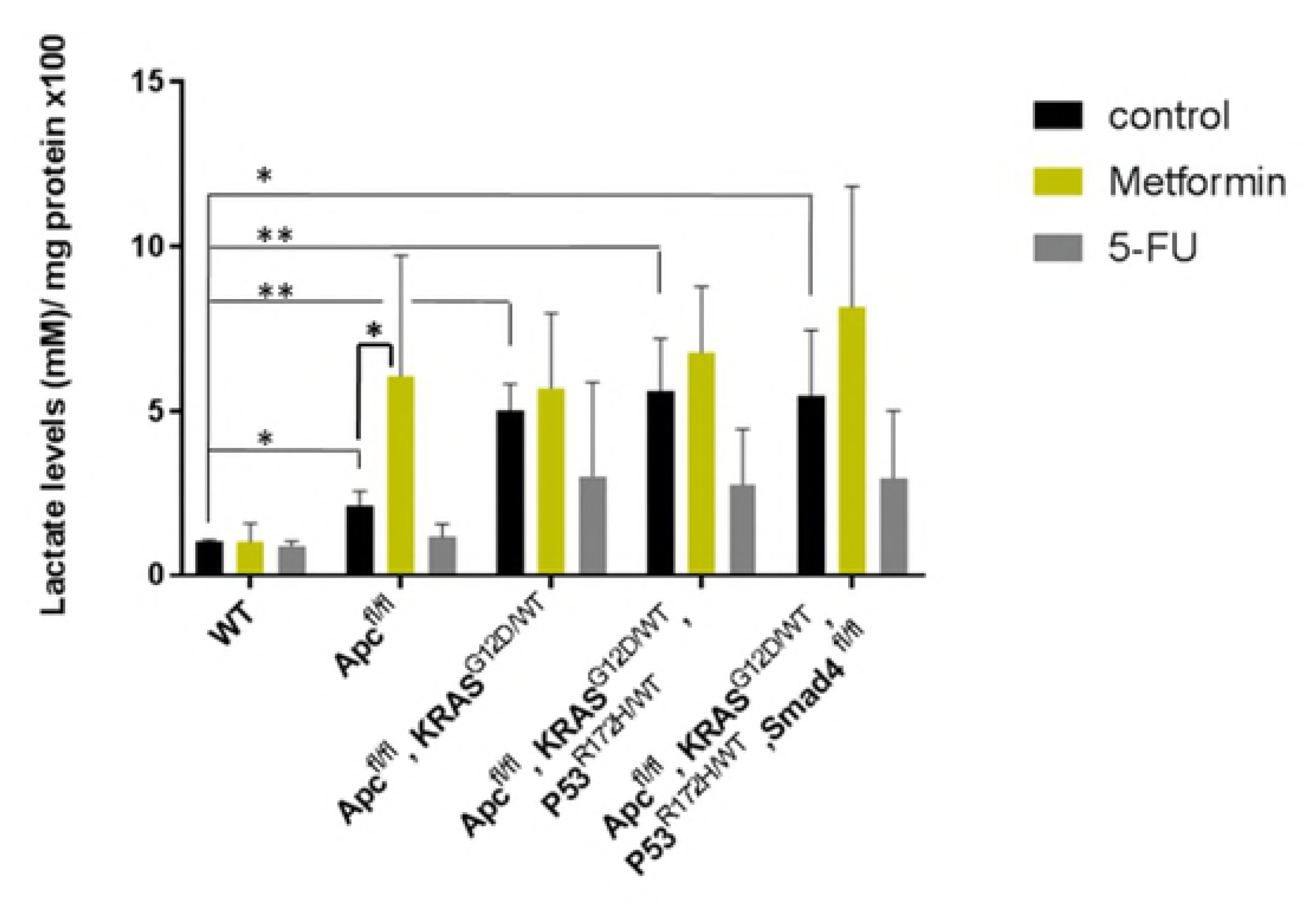
Metformin action in CRC organoids is not related to a Warburg-effect impairement. Bars represent the extracellular L-lactate production upon overnight PBS treatment (black bars), Metformin treatment (yellow bars) and 5-FU treatment (grey bars) using the Cayman’s Glycolysis cell-based assay. L-lactate production measurement is normalized by total protein content (x100). Data are represented by the fold-change mean ±SD (*n* = 3) in all the plots. (ns, *P*> 0.05; *, *P* ≤ 0.05; **, *P* ≤ 0.01; ***, *P* ≤ 0.001; ****, *P* ≤ 0.0001).

## Discussion

Organoids seem to represent a good tool to study lipid metabolism [22] and previous studies employing intestinal organoids have linked the critical role of fatty acid metabolism to the intestinal epithelial integrity *in vivo* [23]. Therefore, we propose this system to get insight into cancer progression mechanisms in regards to fatty acid metabolism and therefore, to assay ACSL/SCD protumorigenic axis action in CRC.

We showed that ACSL4 augmented while miR-19b-1-3p diminished its expression, both progressively, in murine CRC-like organoids. Metformin action compared to the chemotherapeutic agent 5-FU, in terms of viability reduction, was similar; although no significant reduction was found in WT organoids viability with any treatment. Stage I organoids were the most susceptible to Metformin action compared to 5-FU; while further stages presented similar or stronger sensitivity to 5-FU, including WT organoids. Besides, Metformin was able to reduce the intestinal crypt stem cell marker LGR5 in all the stages, together with two other Wnt downstream targets, Ctnnb-1 and Axin-2. Finally, we showed that even though the CRC-organoids series present a growing Warburg effect through the stages consistent with increased L-lactate levels; Metformin action on CRC organoids viability was not related to an ablation of Warburg effect.

The individual role of ACSL isoform 1 [24,25] and 4 [24,26] as well as SCD [27–31] in CRC has been extensively reported. Surprisingly, while ACSL4 mRNA levels are clearly increased through the stages in this organoids model, this was not the case for ACSL1, and SCD was only overexpressed in advanced stages. These results differ from previous ones using human CRC cells which can be due to differential expression in murine tissues compared to human 2D cultures [5] [8]. Nevertheless, the use of murine organoids allows their genetic engineering and to accurately control the mutations for a better mechanistic characterization, rather than patient tumor-derived biopsies with the high variability that each tumor represent. Thus, our CRC model mimics a sporadic colorectal tumor with the common mutations acquired during the progression of this cancer. Due to the organoids results, the overexpression of the three enzymes could be only present in some punctual tumors. However, the overexpression of ACSL4 is preserved in murine organoids with the acquired CRC most common mutations (Fig 1A), indicating a predominant role of this ACSL/SCD component in these cancer progression aspects. The ACSL4 mRNA huge range of expression considering the most mutated stages (Fig 1A) could be explained since stages III and IV in real tumors present an uncontrolled genetic variability with the accumulation of other undetermined mutations. Organoids would be mimicking these uncontrolled stages, compared to the homogeneity presented in 2D cultures. Conversely, ACSL1 static role (S1A Fig) could be due to a lesser implication in tumor development in this system which can be also explained by the fact that the rodent protein is one residue longer (699 amino acids) than the human protein (698 amino acids), making it necessary to study the extent of this dissimilarity. For its part, SCD overexpression has been mainly reported in mesenchymal tissues, rather than epithelial ones, which are the only scaffold for organoids [32,33] giving a reason for the distinctive results found in these epithelial systems among the first stages (S1B Fig).

Regarding miRs expression, miR-19b-1-3p kept its tumor-suppressor role in murine CRC-like organoids, also reported as a good prognosis miRNA, able to target the axis [8]. The immature isoform of miR-19b-1-3p, miR-19b, and other members of the miR-17-92 cluster, where this miRNA is involved, regulate the self-renewal ability of gastric cancer stem cells [34]. The miR-17-92 cluster role is controversial and dependent on the cancer type [35,36]. However, it is interesting the reported role of this miRNA in digestive cancer stem cells, and its role in CRC stem cells may be a potential line of research henceforth. In line with our results, miR-19b was also reported to downregulate suppressor of cytokine signaling 3 (SOC3), modulating chemokine production in intestinal epithelial cells and thereby avoiding intestinal inflammation in Crohn’s disease, which may ultimately prevent the derived disease, CRC [37].

Since Metformin was able to revert the ACSL/SCD EMT phenotype, we tried to gain insight on this process using organoid cultures resembling the different stages of a CRC progression. Metformin treatment seems to be more efficient than 5-FU only during first tumor stages, making organoids recovery harder compared to the ones treated with 5-FU. We propose that Metformin therapies could be an appealing alternative in those cases when the tumor is detected in very early stages rather than 5-FU treatments. However, some studies of Metformin treatment in CRC patients points to stage III to be the most likeable to present an effect [38]. Since CRC is very improbable to detect on its very early stages, known as one of the most silent and deadly cancer; we wonder whether these studies with a low number of candidates in stage I are enough representative.

As well, Metformin therapies has been proposed alone or in combination with other drugs, in CRC. For example, Metformin has been recently combined with aspirin to treat middle stages in non-diabetic CRC patients.( II and III stages) [39]. Furthermore, it exists a Phase 2 Trial for the study of Metformin and 5-Fluorouracil combination in metastatic CRC [40], concluded with a longstanding cancer control. An older report also claimed the benefits of this combination, but they also reported that Metformin alone has antineoplastic activity *per se* in colon cancer cells, and enhanced the activity of 5-FU, oxaliplatin and irinotecan in cells previously treated [41].

Previous reports hypothesized that the inhibition of mitochondrial complex I was the main mechanism of action for Metformin. However, recent studies suggest that cancer progression is compromised upon Metformin treatment, by decreasing the TCA cycle’s anaplerosis. Metformin decreases the flow of glucose- and glutamine-derived metabolic intermediates into the TCA cycle, decreasing the citrate output of the mitochondria and leading to a reduction of acetyl-CoA (Ac-CoA) and oxaloacetate (OAA) in the cytoplasm and therefore a reduction in de novo FA synthesis [42]. This way, Metformin could be targeting lipid metabolism through ACSL/SCD axis. ACSL4 downregulation in the presence of Metformin is clearly evident and the results are larger significant in first stages (I, II) (Fig 4A). Maybe, the reduced overexpression of ACSL4 in the first stages (Fig 1A) increases the sensitivity to Metformin action (Fig 4A); while in more advanced stages, the overexpression is so high that Metformin action could appear to be less effective. This would not be the case for SCD, which showed no overexpression in the first stages and enhanced overexpression in III and IV stages, though it is significantly reduced upon Metformin exposure again in the first stages (Fig 4B). On the other hand, it has been reported that variations in the types and amounts of fatty acids, are able to modify intracellular ACSLs expression [43], thus, this conditions could be also affecting ACSL/SCD components expression besides that the network connection between those enzymes could make them present coordinated effects upon Metformin treatment, reducing its expression due to the lack of their substrate. Metformin was also previously reported to downregulate ACSL expression, lowering fatty acid synthesis and normalizing lipid profile in diabetic rats [44]; as well as limiting its products, 18- carbon chain length fatty acids, in skeletal muscle insulin resistant rats [45], suggesting in this case that metformin is increasing FAs mitochondrial channeling due to the reduction of CPT1 inhibition by malonyl-CoA and therefore decreasing 18-carbon acyl-chain-derived bioactive lipids in the cytoplasm [45], This action of Metformin could be additional to the aforementioned, detoxifying ACSLs probable over activity.

Metformin seems also to target cancer stem cells of different cancer types [46]. However, we have described for the first time the LGR5 downregulation in CRC-like organoids upon Metformin treatment; consistent with previous reports using 2D CRC cultures [47]. LGR5 was diminished in the whole CRC-like organoids series to minimum levels, an indicative that Metformin action is affecting the stem cells of the crypt, responsible for the progression of the organoids lineage. Curiously, Metformin treated organoids do not present apoptosis or even necrosis, but they kept at a minimum size compared to other treatments, where the organoids layer disappeared and the cells appeared apoptotic in the lumen (S6 Fig), showing that cell membrane biogenesis is somehow blocked, mostly built by *de novo* lipogenesis routes.

Finally, Metformin treated CRC organoids exhibit a greater compensatory increase in aerobic glycolysis. Since ATP levels are diminished due to complex I inhibition, the metabolic sensor AMPK is activated, inhibiting mTOR and proliferative events; and promoting glycolysis as an alternative ATP source [48]. We found that even though the CRC-organoids serie presented an increasing glycolysis with the stages, (Fig 5); Metformin was able to increase more this glycolytic phenotype, especially in stage I organoids, coincident with the higher sensitivity to the drug in this organoids. These results point towards Metformin targeting different metabolic routes other that Warburg effect to perform its effect on CRC organoids viability.

Even though the Warburg effect is a priority for current drugs, each day the evidence grows that other metabolic pathways should be targeted for cancer progression ablation. CRC is a leading cause of death in the developed world, though yet simplistic preclinical models that mimic the usual stages of CRC progression are lacking [13]. In this way, organoids further analysis need to be included as the tool of choice for stage-dependent drugs screening.

## Conclusions

### General conclusion

1. Organoids display a precise platform to assay **tumor stage-dependent drugs** being suitable for **personalized medicine**, constituting an invaluable tool due to their relatively low costs, animal saving suffering and their ease and legibility to genetically manipulate.

### Metformin-related conclusions

2. **Metformin** treatment is further proved as an efficient drug in CRC:

- It is able to decrease CRC-like organoids viability at the same rate as current chemotherapy (5-FU) but it does not affect to WT organoids.
- Metformin treatment recovery is significantly inferior compared to 5-FU in **first stages organoids**, but with a greater recovery in WT organoids; becoming an appealing chemotherapy drug in first tumor phases.
- Metformin downregulates the stem cell biomarker LGR5 and Wnt target genes expression in all CRC-like organoid stages, reaffirming its potential use in intestinal cancers.
- Metformin action in CRC organoids is not related to a Warburg-effect impairment, presuming that other metabolisms rather than Warburg should be targeted to complete the cancer progression obstruction

### ACSL/SCD-related conclusions

3. **ACSL4** is progressively overexpressed throughout CRC-like organoids stages; while **miR-19b-1-3p** preserves its protective role, reflecting the role of ACSL/SCD axis action on CRC progression. Besides, Metformin action is stronger on ACSL4 and SCD-overexpressing first stages organoids, agreeing with Metformin greater action on this stage.

## Acknowledgment

We kindly thank Dr. Daniel Stange and Dr. Schölch for providing the CRC-like organoids and to the whole Department of Gastrointestinal, Thoracic and Vascular Surgery at University Hospital Carl Gustav Carus for its continuous support. The research stay on this lab was kindly financed with the assistance of a Boehringer Ingelheim Fonds travel grant.

## Conflict of interest statement

Authors declare no potential conflict of interest.

## Source of funding

This work was supported by Ministerio de Economía y Competitividad del Gobierno de España (MINECO, Plan Nacional I+D+i AGL2016-76736-C3), Gobierno regional de la Comunidad de Madrid (P2013/ABI-2728, ALIBIRD-CM) and EU Structural Funds.

## Supporting information

**S1 Table**. Primers’ sequences (Invivogen) used for quantitative real-time PCR.

**S2 Table**. Probes from TaqMan^®^ MicroRNA Assays (ThermoFisher) used for quantitative real-time PCR.

**S1 Fig. ACSL1 and SCD mRNA expression throughout CRC-like organoids stages**

RT-QPCR analysis showing ACSL1 (A) and SCD (B) mRNA expression levels throughout CRC-like organoids stages. Results represent the fold-change mean ±SD (*n* = 3) (ns, *P* > 0.05; *, *P* ≤ 0.05; **, *P* ≤ 0.01; ***, *P* ≤ 0.001; ****, *P* ≤ 0.0001)

**S2 Fig. ACSL/SCD regulatory miRNAs expression in CRC-like organoids**

RT-QPCR analysis showing mRNA expression levels throughout CRC-like organoids stages of different ACSL/SCD regulatory miRNAS: miR-19b-1-5p (A), miR-142-3p (B), miR-142-5p (C). Results represent the fold-change mean ±SD (*n* = 3). (ns, *P*> 0.05; *, *P* ≤ 0.05; **, *P* ≤ 0.01; ***, *P* ≤ 0.001; ****, *P* ≤ 0.0001).

**S3 Fig. Metformin and 5-FU effect in CRC-like organoids.**

MTT cell viability assays upon 48 hours treatments with PBS (black bars), 10 uM Metformin (yellow bars) or 10, 100 and 150 uM 5-FU (grey bars) in the different CRC-like organoids representative stages: WT organoids (A); *APC*^fl/fl^ organoids resembling stage I (B); *APC*^fl/fl^ *, KRAS*^G12D/WT^ organoids resembling stage II (C); *APC*^fl/fl^ *, KRAS*^G12D/WT^*, P53*^R172H/WT^ organoids resembling stage III (D); *APC*^fl/fl^ *, KRAS*^G12D/WT^ *,P53*^R172H/WT^, Smad4^fl/fl^ organoids resembling stage IV (E). Data are represented by the fold-change mean ±SD (*n* = 3) in all the plots. (ns, *P* > 0.05; *, *P* ≤ 0.05; **, *P* ≤ 0.01; ***, *P* ≤ 0.001; ****, *P* ≤ 0.0001).

**S4 Fig. Metformin and 5-FU recovery effect in CRC-like organoids.**

MTT cell viability assays upon 48 hours treatments (black bars) and upon extra 72h post-treatment recovery with PBS (light grey bars), 10 uM Metformin (yellow bars) or 10, 100 and 150 uM 5-FU (dark grey bars) in the different CRC-like organoids representative stages:: WT organoids (A); *APC*^fl/fl^ organoids resembling stage I (B); *APC*^fl/fl^ *, KRAS*^G12D/WT^ organoids resembling stage II (C); *APC*^fl/fl^ *, KRAS*^G12D/WT^ *,P53*^fl/R172H^ organoids resembling stage III (D); *APC*^fl/fl^ *, KRAS*^G12D/WT^ *,P53*^fl/R172H^,Smad4^fl/fl^ organoids resembling stage IV (E). Data are represented by the fold-change mean ±SD (*n* =3) in all the plots. (ns, *P* > 0.05; *, *P* ≤ 0.05; **, *P* ≤ 0.01; ***, *P* ≤ 0.001; ****, *P* ≤ 0.0001).

**S5 Fig. ACSL/SCD axis and stem cell markers expression (Lgr5, Axin-2 and Ctnnb-1) upon Metformin and 5-FU treatment**

Expression leveles of enzymes related to the ACSL/SCD axis, ACSL1 (A), ACSL4 (B) and SCD (C) by RT-QPCR; and expression levels of different stem cell markers, Lgr5 (D), Axin-2 (E) and Ctnnb-1 (F) by RT-QPCR; upon PBS (black bars), 10 uM Metformin (yellow bars) or 10, 100 and 150 uM 5-FU (grey bars). Data are represented by the fold-change mean ±SD (*n* = 3). (ns, *P*> 0.05; *, *P* ≤ 0.05; **, *P* ≤ 0.01; ***, *P* ≤ 0.001; ****, *P* ≤ 0.0001).

**S6 Fig. Comparative organoids morphology between Metformin and other oncologic treatments.**

Organoids (stage I and III) representative pictures with DMSO, Metformin and other metabolic drugs against CRC progression, upon 48 hours treatments plus upon extra 72h post-treatment recovery. Pictures were captured using the × 10 objective, in bright field. Leica microscope (Leica microsystems).

## Abbreviations

ACSL1: Acyl-CoA synthetase 1
ACSL4: Acyl-CoA synthetase 4
CRC: Colorectal cancer
EMT: Epithelial-mesenchymal transition
5-FU: Fluorouracil
MiRNAs/ miR: MicroRNAs
MTT: 3-(4, 5-dimethylthiazol-2-yl)-2, 5-diphenyltetrazolium bromide
OAA: Oxaloacetate
SCD: Stearoyl-CoA desaturase
TCA: Tricarboxylic Acid cycle
2D: 2-dimensional
3D: 3-dimensional.

